# Structural pre-training improves physical accuracy of antibody structure prediction using deep learning

**DOI:** 10.1101/2022.12.06.519288

**Authors:** Jarosław Kończak, Bartosz Janusz, Jakub Młokosiewicz, Tadeusz Satława, Sonia Wróbel, Paweł Dudzic, Konrad Krawczyk

**Affiliations:** NaturalAntibody, Kraków, Poland

## Abstract

Protein folding problem obtained a practical solution recently, owing to advances in deep learning. There are classes of proteins though, such as antibodies, that are structurally unique, where the general solution still lacks. In particular, the prediction of the CDR-H3 loop, which is an instrumental part of an antibody in its antigen recognition abilities, remains a challenge. Antibody-specific deep learning frameworks were proposed to tackle this problem noting great progress, both on accuracy and speed fronts. Oftentimes though, the original networks produce physically implausible bond geometries that then need to undergo a time-consuming energy minimization process. Here we hypothesized that pre-training the network on a large, augmented set of models with correct physical geometries, rather than a small set of real antibody X-ray structures, would allow the network to learn better bond geometries. We show that fine-tuning such a pre-trained network on a task of shape prediction on real X-ray structures improves the number of correct peptide bond distances. We further demonstrate that pre-training allows the network to produce physically plausible shapes on an artificial set of CDR-H3s, showing the ability to generalize to the vast antibody sequence space. We hope that our strategy will benefit the development of deep learning antibody models that rapidly generate physically plausible geometries, without the burden of time-consuming energy minimization.

## Introduction

Antibodies are the largest group of biologics ^1^, recently reaching hundred approved molecules ^2^ with more forecasted to come to clinical use ^3^. Because of their therapeutic role, there is much interest in the development of computational methods addressing these molecules ^4^.

One of the crucial properties of an antibody to be predicted is its structure ^5^. The antigen-recognizing variable domain has an immunoglobulin fold, with a largely conserved framework region and three complementarity-determining region loops (CDRs) defining the antigen-recognizing paratope. Of the three loops the third CDR of the heavy chain (CDR-H3) is particularly problematic for structure prediction ^6–9^. Methods such as AlphaFold2 ^10^ have made great progress in general protein structure prediction, however, some of its applications lack in face of antibodies whose CDRs undergo a different evolutionary process (evolving in days rather than millions of years), and thus are structurally unique ^11^.

To address this issue, but also to draw from AlphaFold2 and its derivatives ^12,13^, many antibody-specific deep learning structure prediction methods have been introduced ^4,13^. The algorithms started by addressing CDR-H3 loop prediction such as DeepH3 ^7^ and AbLooper ^8^, after extending these to the entire variable domain via NanoNet ^14^ and the entire Fv molecule by DeepAb ^15^, IgFold ^16^, AbodyBuilder2 ^17^, EquiFold ^18^ and tFold-Ab ^19^. As opposed to the homology methods that reported CDR-H3 root mean squared deviation (RMSD) accuracies in the region of 3-4Å ^9^, the deep learning methods achieve an RMSD of 2-3Å RMSD. The methods are also faster -achieving predictions in seconds, rather than in minutes, as was the case with best homology modeling tools ^20,21^ and tens of minutes in case of AlphaFold2.

The methods introduced a variety of architectures such as residual convolutional networks ^15^, graph neural networks ^8^ or drawing from the structure of AlphaFold2 ^16,17^. In certain guises, large-scale sequence information from the Observed Antibody Space ^22,23^ was introduced to provide a sequence pre-training basis for the models ^16^. The methods achieve similar accuracies to each other and the original AlphaFold, perhaps with a minor advantage from focusing on the antibody datasets.

Though in statistical terms the methods do not provide an order of magnitude improvement in antibody prediction accuracy with respect to AlphaFold2, they have a great advantage in terms of faster running time. Coordinates can be generated in seconds, rather than in minutes or even hours as is the case with AlphaFold2. Though the networks produce the results fast, they are afflicted by problems relating to the physical plausibility of the generated structures ^24^. The problem was highlighted by methods such as Ablooper, IgFold, and AbodyBuilder2, as they can produce the coordinates rapidly, but then the final structure needs to be energy minimized ^8,16,25^.

One can address such issues in coordinate generation by employing special loss functions such as the Frame Aligned Point Error (FAPE), or the distogram prediction loss in AlphaFold2 ^10^. One can also introduce physical priors to improve the quality of generated coordinates ^26^. In place of employing explicit energy calculator programs such as OpenMM, one can attempt to train a neural network to approximate such functions ^27^.

Here we address the problem of improving antibody structure prediction by generating more physically plausible structures directly from the network. An important aspect that impacts model performance is the availability of data that can be used in the training process. With antibody diversity estimated to be 10^18^ unique molecules, the publicly available 6,500 of redundant experimentally resolved antibody structures are a relatively small sample. Most of the models that attempted antibody structure prediction were trained on such a small publicly available dataset of antibody structures, and to the best of our knowledge only IgFold employed AlphaFold to augment its input dataset. Such a small dataset of antibodies is clearly powerful enough to teach the broad features of an immunoglobulin shape but falls short of making the structures physically plausible out-of-the-network. To address this issue, one might attempt to augment input dataset to expose the network to larger amount of correct bond geometries. Here we demonstrate that pre-training a network on a large, augmented dataset of antibody model structures allows it to learn such physical features.

## Results

### Problem setup

We test the hypothesis that pre-training on a large set of model structures allows the network to learn physically plausible conformations that can then be fine-tuned on the real dataset of crystal structures. For this effort, we focus on the distances of the peptide bond, simplified to the distance between consecutive Cα carbons. In crystal structures, the distribution of such distances is firmly centered on 3.8Å, with a small standard deviation in the order of 0.1Å. Current deep learning models are trained to have the distribution of Cα distances centered on 3.8, however, with a much larger standard deviation. This results from the models oftentimes producing peptide bonds that are implausibly far away or close together (Figure 1) -though they are close enough in RMSD space to the template that they aim to reproduce.

**Figure 1.**
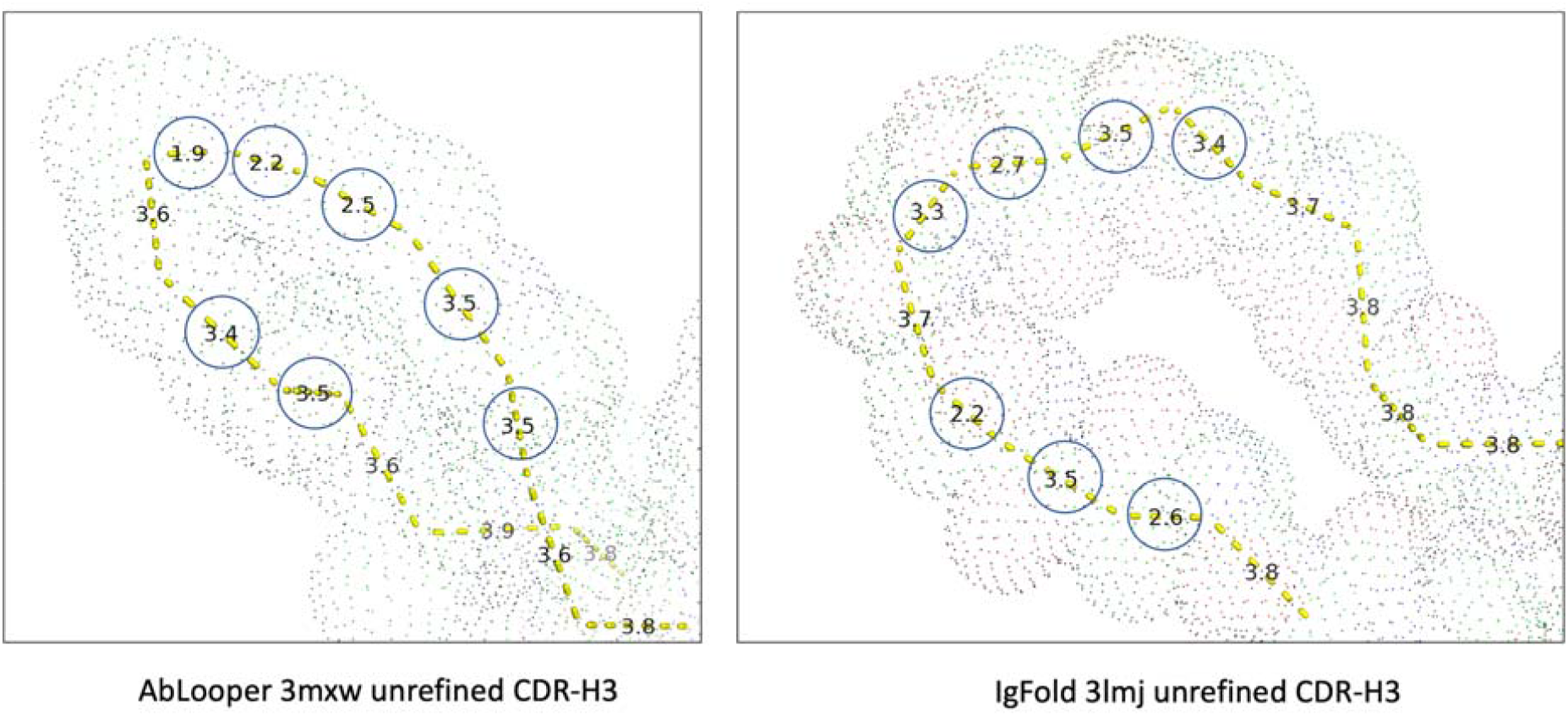
Example of CDR-H3 predictions with consecutive Cα distances requiring refinement. Distances outside of the range of [3.6Å 3.9Å] range are circled. The two structures are outputs from the neural networks of AbLooper and IgFold, without applying refinement post-processing. After energy refinement using OpenMM and Rosetta respectively for each method, the distances become physically plausible, however at a high computational cost.

As the basis architecture, we aimed for the simplest deep learning model that would not rely on improvements such as problem-specific loss functions or all-atom predictions. For this purpose, we re-created the architecture of NanoNet ^14^. The model is a small residual neural network (seven blocks, 1.9m parameters) predicting coordinates directly, in comparison to other networks using more advanced architectures and training regimes. In its published form NanoNet predicts all the backbone atoms, however, we implemented a version only predicting Cα to make sure we provide the model with minimal information.

For training & testing the network, we employed two augmented sets of homology models for pre-training, a set of crystal structures for fine-tuning and the Rosetta test set as the hold-out. The Rosetta test set was previously used to benchmark several models ^7,8^, but we reduced it to 45 structures due issues such as incomplete chains. The crystal structure dataset was obtained from the PDB by identifying antibodies therein ^28,29^. Here we have three datasets, CrystalHeavy, consisting of heavy chain structures, CrystalLight, consisting of light chain structures and CrystalCombined, where we merged the two previous datasets. We used two pre-training datasets, *‘*Natural*’* and *‘*Homology*’*. The Homology dataset results from homology modeling a subset of approximately 40,000 paired structures available in OAS ^22,23^. To control whether the results we obtain are the function of the dataset we chose and the homology method we implemented, we also employed the Natural dataset that was created independently of NaturalAntibody. This consisted of the public-response structures, carefully curated from naturally occurring antibodies in NGS datasets ^30^. We also created a combined pre-training dataset consisting of all heavy and light chains in the Natural and Homology datasets, termed PretrainingCombined. All the datasets employed are briefly described in Table 1, with details in Methods.

**Table 1.**
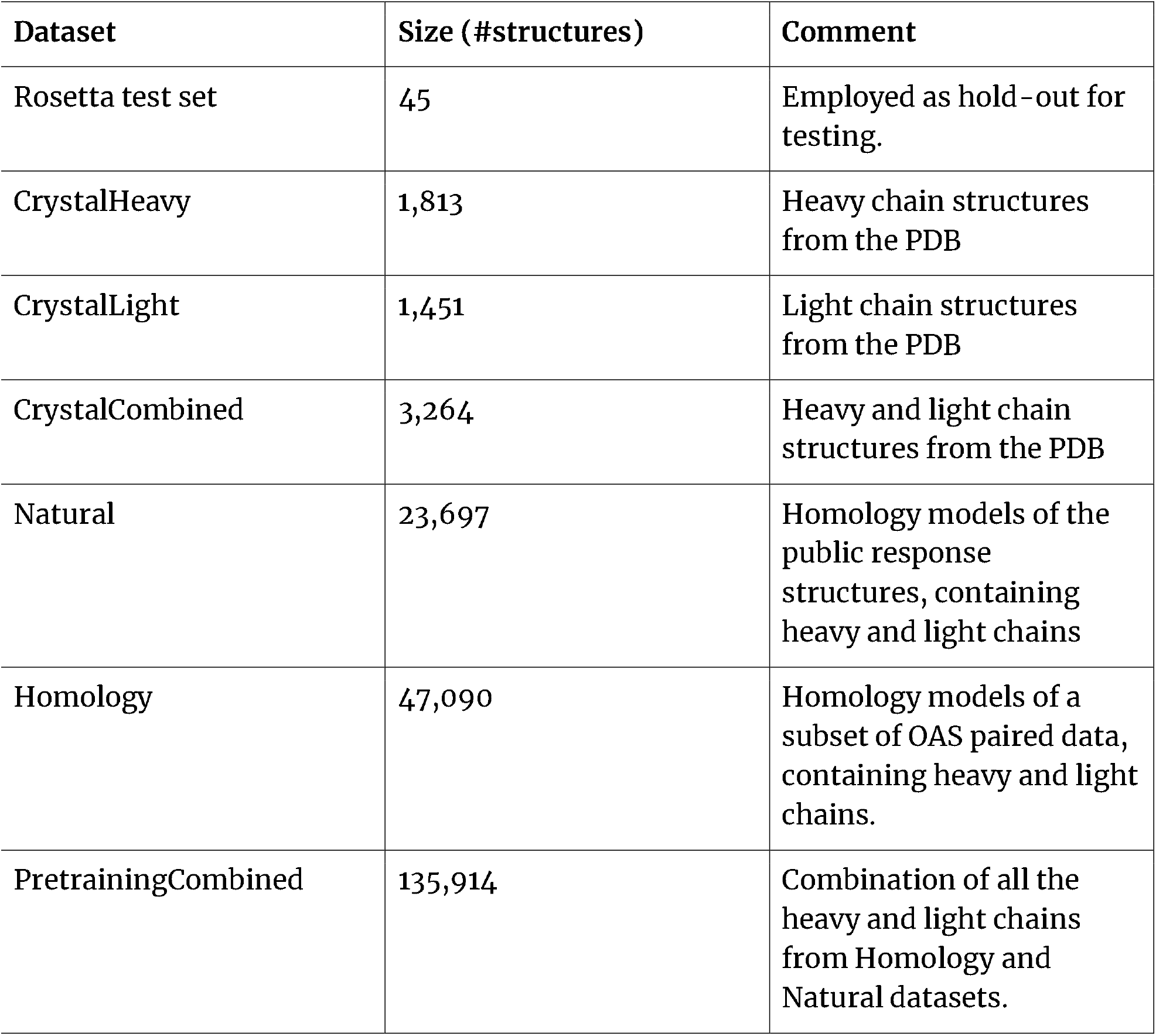
Datasets employed in this study.

| **Dataset** | **Size (#structures)** | **Comment** |
| --- | --- | --- |
| Rosetta test set | 45 | Employed as hold-out for testing. |
| CrystalHeavy | 1,813 | Heavy chain structures from the PDB |
| CrystalLight | 1,451 | Light chain structures from the PDB |
| CrystalCombined | 3,264 | Heavy and light chain structures from the PDB |
| Natural | 23,697 | Homology models of the public response structures, containing heavy and light chains |
| Homology | 47,090 | Homology models of a subset of OAS paired data, containing heavy and light chains. |
| PretrainingCombined | 135,914 | Combination of all the heavy and light chains from Homology and Natural datasets. |

### Structural and sequence similarities between pre-training data and real crystal structures - controlling for data leaks

There exists a risk of data leakage between the pre-training and crystal structure datasets. The similarities could arise from 1) data leakage of templates between pre-training and crystal structure datasets 2) close sequence identity between pre-training and crystal structure datasets as well as 3) close data leakage as a result of test templates being employed in generating the datasets.

In our Homology pre-trained dataset, we ensured that no templates were being used that were in the Rosetta test set, judging by identical CDR-H3/L3 sequences. No such control was possible for the Natural dataset as it was pre-prepared. Attempting a conservative approach, we purged the Homology and Natural datasets from sequences that had a close resemblance to those in the entire crystal structure dataset as judged by identical CDR-H3 or CDR-L3.

Even after removing all the non-identical CDR-H3s and CDR-L3s, the pre-training datasets still had structures within 1Å RMSD of one in the Rosetta training set, which were subsequently removed. A total of 4761 structures were removed from the Homology dataset and 339 from the Natural dataset. Afterwards, the Homology dataset had an average CDR-H3 RMSD of 2.8Å to the closest CDR-H3 in the Rosetta test set with the corresponding figure for the Natural dataset being 3.2Å. In total, 146 structures in the crystal dataset had an RMSD of less than 1Å to the closest one in the Rosetta test set, indicating that diverse sequences of CDR-H3 can still adopt similar folds.

Finally, we also checked the overlap between the structural training set *‘*purged*’* of identical CDR-H3s, with that of the Rosetta test set, given in Table 2. Overall, we did not note radical closeness of the pre-training datasets to that of the Rosetta test set.

**Table 2.** CDR-H3 Sequence overlaps between the CrystalHeavy and the pre-training datasets. The alignments were computed for length-matched IMGT CDR-H3s using the biopython pairwise alignment function.

|  | Rosetta Test Set (n=45) |  |  | Crystal Structures (CrystalHeavy) (n=1,813) |  |  |
| --- | --- | --- | --- | --- | --- | --- |
| | $\mu$ | Max | Min | $\mu$ | Max | Min |
| Crystal Structures (CrystalHeavy) | 57% | 92% | 25% | n/a |  |  |
| Natural pre-training | 54% | 77% | 33% | 46% | 88% | 0% |
| Homology pre-training | 66% | 91% | 38% | 59% | 92% | 0% |

### Effect of structural pre-training on the number of incorrect peptide bonds in heavy chain prediction

Equipped with two pre-training sets, Natural and Homology, we used our interpretation of the NanoNet model to see whether pre-training has any effect on the number of incorrect peptide bonds. We defined the *‘*incorrect*’* peptide bonds as those falling outside of the range [3.6Å,3.9Å].

For training, we employed a simplistic loss function that consisted of the Mean Squared Error in Cα of the predicted vs template positions, together with the consecutive Cα distance loss parameter. The Cα distance loss component is weighted, with higher weights incurring a larger penalty for consecutive Cα*’*s distance being different from 3.8Å. For training, we checked the effect of the Cα loss weight of 1, 5, 10, 15, 20.

For each value of the Cα loss weight, we trained six models to predict the heavy chain. Three models without any benefit of pre-training, each for Natural, Homology, and CrystalHeavy with a learning rate of 10^−3^. Such models were named after the dataset and the Cα weight, e.g. CrystalHeavy model with the Cα weight of 5 would be called CrystalHeavy5. Then, each of the three models served as a pre-training basis for training on the CrystalHeavy structure dataset with a learning rate of 10^−5^. In each case of fine-tuning the Cα weight was the same as in pre-training. A fine-tuned model would be called by combining its pre-trained name with CrystalHeavy and the Cα parameter. For instance, a model pre-trained on Homology with the Cα weight of 10 would be Homology10CrystalHeavy10. Using the Crystal dataset for both pre-training and fine-tuning was supposed to contrast the effects of using the Natural and Homology pre-training datasets. Each of the models was benchmarked on the Rosetta test set, noting the number of peptide bonds outside of the [3.6,3.9] range, median and standard deviation of the peptide bond distribution as well as the Cα RMSD of the CDR-H3 (after aligning the entire V-region). All the results are compiled in Table 3.

**Table 3.** Structural pre-training improves the peptide bond length quality of the model. We checked to what extent increasing the weight on the Cα bond in the loss function improves the matters vs pretraining. In each case, increasing the Cα bond weight in loss reduces the number of bad bonds. However in each case, consistently, the models that had been pre-trained achieved the best results, on both pre-training datasets. For contrast, the very bottom row shows the peptide bond Cα distances from the crystal structures in the Rosetta test dataset. The results in the table are grouped by their Cα loss penalty weights.

| Model | Cα weight | #d < 3.6Å | #d > 3.9Å | Sum bad bonds | $\sigma$ | $\mu$ | Mean IMGT H3 RMSD |
| --- | --- | --- | --- | --- | --- | --- | --- |
| CrystalHeavy1 | 1 | 1330 | 1176 | 2506 | .26 | 3.73 | 2.11 |
| Natural1 | 1 | 665 | 501 | 1166 | .19 | 3.73 | 2.79 |
| Homology1 | 1 | 976 | 372 | 1348 | .19 | 3.70 | 2.06 |
| CrystalHeavy 1<br>CrystalHeavy 1 | 1 | 1091 | 974 | 2065 | .23 | 3.72 | 2.09 |
| Natural1<br>CrystalHeavy 1 | 1 | 477 | 597 | ~~1074~~ | .16 | 3.75 | 2.20 |
| Homology1<br>CrystalHeavy 1 | 1 | 603 | 634 | 1237 | .16 | 3.75 | 2.04 |
| CrystalHeavy 5 | 5 | 328 | 978 | 1306 | .13 | 3.79 | 2.15 |
| Natural5 | 5 | 426 | 843 | 1269 | .16 | 2.77 | 2.26 |
| Homology5 | 5 | 702 | 529 | 1231 | .18 | 3.72 | 2.02 |
| CrystalHeavy 5<br>CrystalHeavy 5 | 5 | 503 | 493 | 996 | .12 | 3.75 | 2.17 |
| Natural5<br>CrystalHeavy 5 | 5 | 208 | 641 | 849 | .11 | 3.79 | 2.16 |
| Homology5<br>CrystalHeavy 5 | 5 | 209 | 563 | ~~772~~ | .11 | 3.78 | 2.10 |
| CrystalHeavy 10 | 10 | 345 | 546 | 891 | .11 | 3.76 | 2.19 |
| Natural10 | 10 | 334 | 352 | 686 | .13 | 3.75 | 2.26 |
| Homology10 | 10 | 238 | 741 | 979 | .12 | 3.78 | 2.14 |
| CrystalHeavy 10<br>CrystalHeavy 10 | 10 | 300 | 396 | 696 | .10 | 3.75 | 2.20 |
| Natural10<br>CrystalHeavy 10 | 10 | 117 | 316 | 433 | .08 | 3.78 | 2.21 |
| Homology10Crystal<br>Heavy10 | 10 | 182 | 238 | <u>420</u> | .08 | 3.76 | 2.22 |
| CrystalHeavy15 | 15 | 272 | 835 | 1107 | .14 | 3.78 | 2.10 |
| Natural15 | 15 | 296 | 444 | 740 | .11 | 3.77 | 2.34 |
| Homology15 | 15 | 310 | 292 | 602 | .13 | 3.75 | 2.20 |
| CrystalHeavy15Cryst<br>alHeavy15 | 15 | 135 | 408 | 543 | .9 | 3.78 | 2.10 |
| Natural15CrystalHea<br>vy15 | 15 | 126 | 362 | 488 | .8 | 3.78 | 2.27 |
| Homology15Crystal<br>Heavy15 | 15 | 122 | 189 | <u>311</u> | .7 | 3.77 | 2.16 |
| CrystalHeavy20 | 20 | 265 | 600 | 865 | .13 | 3.77 | 2.19 |
| Natural20 | 20 | 271 | 185 | 456 | .10 | 3.75 | 2.19 |
| Homology20 | 20 | 298 | 394 | 692 | .12 | 3.76 | 2.25 |
| CrystalHeavy20Cryst<br>alHeavy20 | 20 | 111 | 396 | 507 | .08 | 3.78 | 2.26 |
| Natural20CrystalHea<br>vy20 | 20 | 83 | 358 | 441 | .07 | 3.79 | 2.19 |
| Homology20Crystal<br>Heavy20 | 20 | 50 | 349 | <u>399</u> | .07 | 3.79 | 2.14 |
| **Rosetta test set**<br>**crystal structures** | n/a | 1 | 25 | 26 | .05 | 3.8 | n/a |

Though there are discrepancies between CDR-H3 modeling quality, all the models appear to end up within a quality range, rather than specific models being better than others by an order of magnitude.

In each case reported in Table 3, models that had the benefit of pre-training using either the Homology or Natural dataset produced better peptide bond lengths than base models without pre-training. In contrast, pre-training on the CrystalHeavy structure dataset led to worse peptide bond lengths than pre-training on either Natural or Homology dataset, showing the benefit of the different base models. The number of incorrect peptide bonds is lower with pre-training without a significant effect on the quality of CDR-H3 prediction.

As expected, the higher the weight for the Cα penalty, the lower the number of peptide bonds falling outside of our predefined erroneous range. Note, however, that even the highest values for the loss penalty we employed in Table 3 (20), did not approach the crystal structure number of bad bonds on the same dataset which is 26. To test the limits of the Cα penalty, we employed a radically higher weight of 100. The resulting CrystalHeavy100 model achieved a radically worse number of peptide bonds, than models with lower penalties (3,910). By contrast the Natural100Crystal100 model achieved 138 incorrect bonds. This suggests that there are limits to biasing the loss function that can be overcome by the benefit of a larger pre-training test set.

Altogether, these results demonstrate that using structural pre-training, appears to reduce the number of incorrect peptide bonds independently of the weight of Cα and the training regime, without the detriment to the CDR-H3 RMSD accuracy.

### Effect of mixing heavy and light chain predictions

The previous experiment consisted entirely of the heavy chain predictions. We, therefore, tested a hypothesis that creating a combined predictor of heavy and light chains would offer better results on the constraints of peptide bonds, by the same virtue of using a larger dataset.

We employed three training datasets/regimes -heavy chain only (CrystalHeavy), light chain only (CrystalLight), and combining the heavy and light prediction (CrystalCombined). Each model was trained in three copies to test the reproducibility. We employed the Cα penalty of 3.0, so as not to give it too much bias but also so as not to leave the model too reliant on the template MSE loss as well. The heavy models were tested on the CDR-H3 structures from the Rosetta test, the light ones on the CDR-L3 from the Rosetta test, and the combined set on both. The results are compiled in Table 4.

**Table 4.**
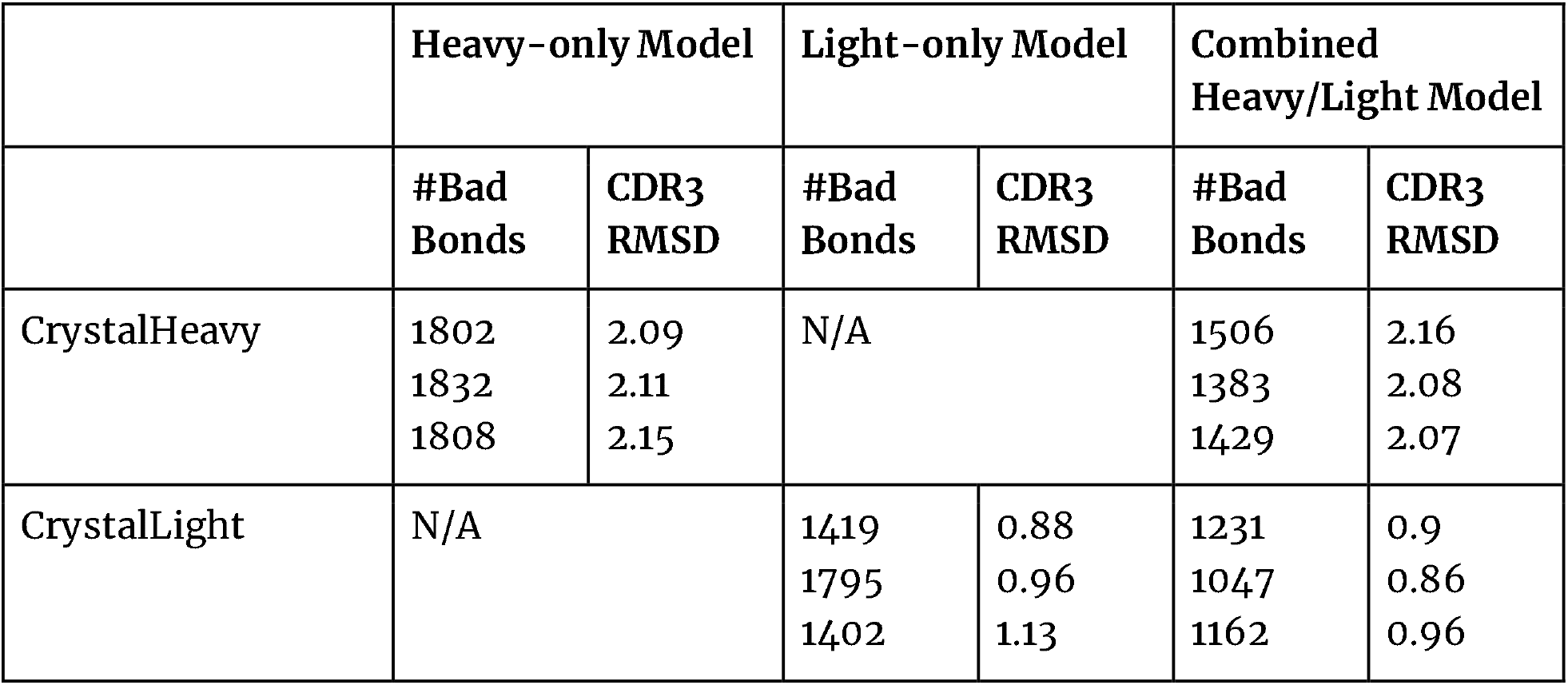
Contrasting of CDR-H3/CDR-L3 modeling between chain-specific models and the combined model. We trained three different models on a different train/validation split on the CrystalHeavy, CrystalLight and CrystalCombined datasets. We used the Rosetta test set to verify the CDR-H3 and CDR-L3 predictions only. It appears that combining heavy and light chain predictions does not improve the accuracy of CDR3 prediction based on RMSD. However, it appears to contribute to a smaller number of bad peptide bonds (<3.6Å or >3.9Å) in the resulting structures.

Judging by RMSD accuracy in Table 4, predicting the CDR-L3 chain is a much easier task for the network than CDR-H3 prediction -except for 3mlr which is an outlier with a shifted beta sheet (Figure 2). We do not note any observable benefit or detriment to the prediction of CDR-H3 or CDR-L3 by combining the predictions. However, it appears that having access to more structures allowed the combined model to reduce the number of incorrect peptide bonds across heavy and light chains. This provides supplementary evidence that augmenting the datasets in antibody modeling appears to have a favorable effect on the basic physical plausibility of produced structures without significant detriment, nor benefit to the quality of the estimated backbone shape.

**Figure 2.**
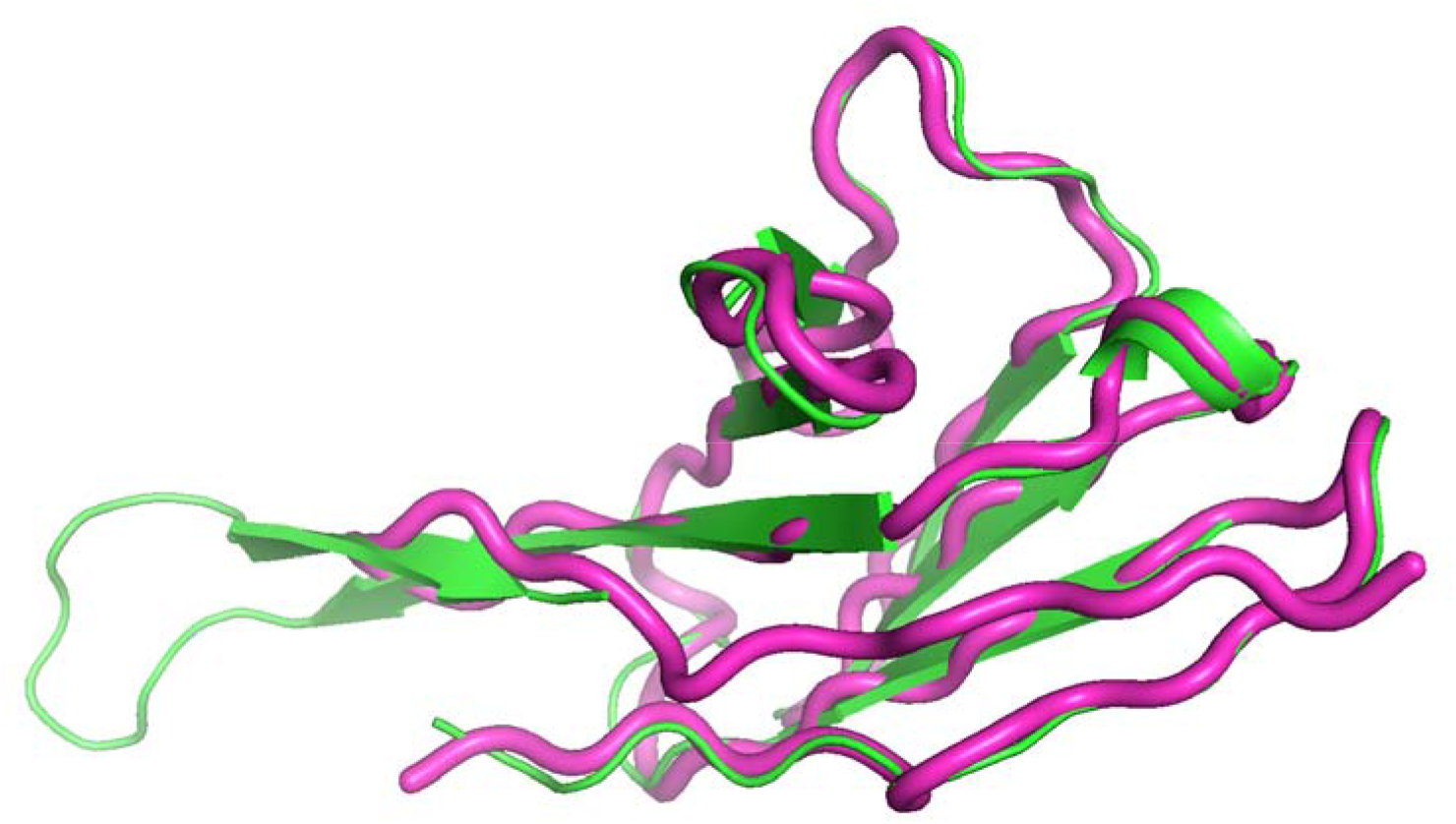
3mlr outlier in light chain modeling. Our prediction (pink) has been overlaid on the native structure (green). One beta strand appears to shift with respect to where it would normally be expected in the immunoglobulin fold. Our modeling consistently failed in that case and produced CDR-L3 RMSD in the region of 10Å.

### Benchmarking antibody structure prediction methods for their ability to produce plausible physical peptide bond lengths

We aimed to establish how the effects of CDR-H3 and peptide bond length accuracies are placed in the context of other available methods. For this reason, we tested those that were freely available for download & academic purposes. These criteria were satisfied by AbLooper, IgFold, AbodyBuilder2, and ESMFold.

All the methods except for AbLooper produce full antibody structures. Ablooper requires the framework to be modeled and predicts the CDRs only. We supplied the real crystal structures of our Rosetta test set as a basis for AbLooper. IgFold, AbodyBuilder2, and AbLooper produce excellent peptide bond lengths when allowed the refinement step of the structure produced by the network. Therefore, for the analysis here, we employed the unrefined structures produced by these methods straight from the network. The ESMFold models were used directly as produced by the network.

Given the findings from previous sections, we created a combined heavy/light chain pre-trained model, by merging the Homology and Natural datasets together (PretrainedCombined). We employed Cα loss penalty of 100 and thus called the model PretrainedCombined100. This model was designed to be a radical version that reduces the number of bad peptide bonds. For contrast we also employed CrystalHeavy1 and CrystalHeavy100 models, to demonstrate the effects of no-pretraining and high Cα penalty loss.

We only focused on the prediction of Cα CDR-H3 accuracy, as the methods produce different outputs from the network - AbLooper only does CDRs, NanoNet and IgFold only backbone and ESMFold and AbodyBuilder2 produce side-chains as well. The CDR-H3 predictions are aligned to the original crystal structures and the RMSDs for each method are given in Figure 3. Even though RosettaAntibody was not a test set for all methods, (validation set for AbodyBuilder2 and no control for ESMFold), all the methods appear to produce a similar distribution of their predictions (Figure 3). This could be indicative of the limits of current deep learning methods in backbone shape prediction. This is of course notwithstanding producing supplementary output, such as additional backbone atoms to Cα, side chains, or the physical plausibility of the structures.

**Figure 3.**
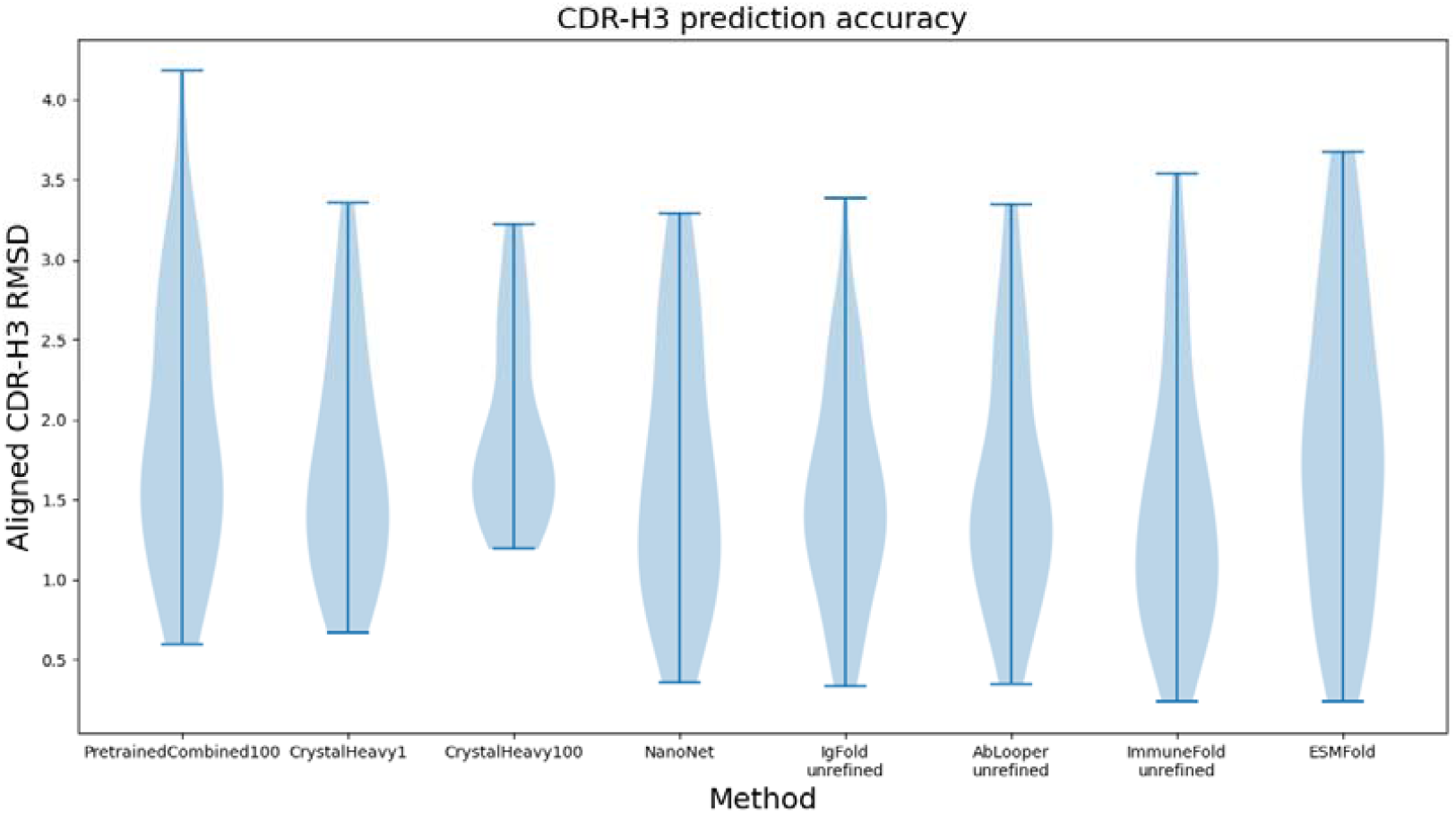
Accuracy of the benchmarked models on the Rosetta dataset. The CDR-H3 were structurally aligned and consecutive Cα distances and RMSD were calculated. The Rosetta test set was also a test set for IgFold and AbLooper. It was a validation set in the case of AbodyBuilder2 and there was no control possible in the case of ESMFold. Despite no proper control for test in such cases, all the methods appear to be in a similar range for their predictions, suggesting that the deep learning predictions of CDR-H3 converge in their accuracy. The error of such CDR-H3 prediction was calculated to be 0.25Å, as the RMSD of H3s of the same length resolved multiple times.

In Table 5 we summarize the number of bad peptide bond distances according to our definition that all the methods in Figure 3 achieved. Unlike in Table 3 where we calculated the total number of bad Cα distances, here we only calculated the peptide bonds in the IMGT-Defined CDR-H3 region. Our pre-trained network achieved the least number of bad peptide bond distances, followed by ESMFold and IgFold. The training regimen that did benefit from pre-training (CrystalHeavy100), obtained a notably worse number of bad peptide bond distances, 527 vs 20 in PretrainedCombined100. Furthermore, the predictions of the PretrainedCombined100 are well centered on 3.8Å (Figure 4), with similar distribution shape in ESMFold. Other methods appear to produce distances even in the range below 3Å.

**Table 5.**
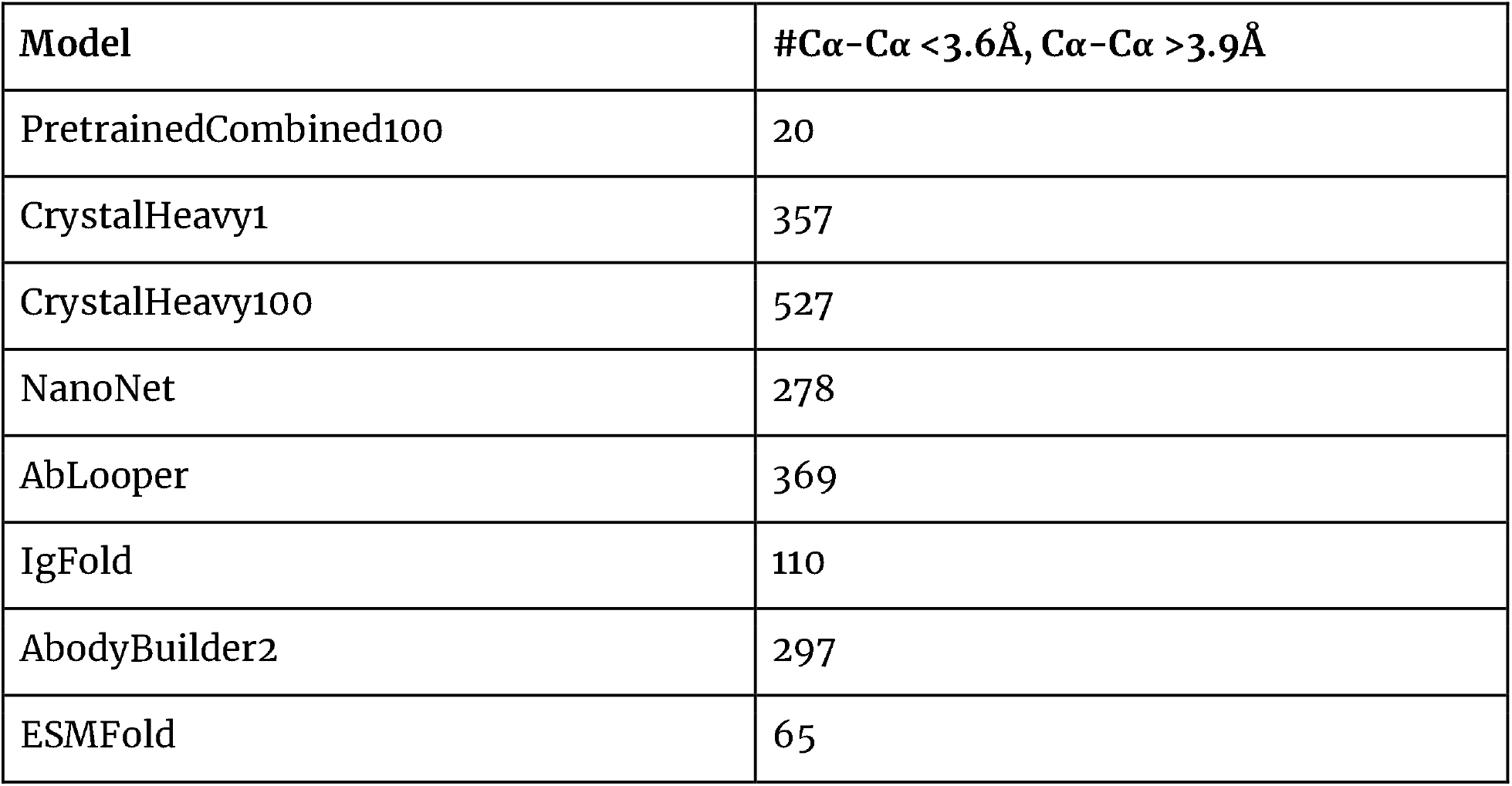
Number of bad peptide bonds on the Rosetta test benchmark. The peptide bond distances were calculated on IMGT-defined CDR-H3 only.

| Model | #Cα-Cα < 3.6Å, Cα-Cα > 3.9Å |
| --- | --- |
| PretrainedCombined100 | 20 |
| CrystalHeavy1 | 357 |
| CrystalHeavy100 | 527 |
| NanoNet | 278 |
| AbLooper | 369 |
| IgFold | 110 |
| AbodyBuilder2 | 297 |
| ESMFold | 65 |

**Figure 4.**
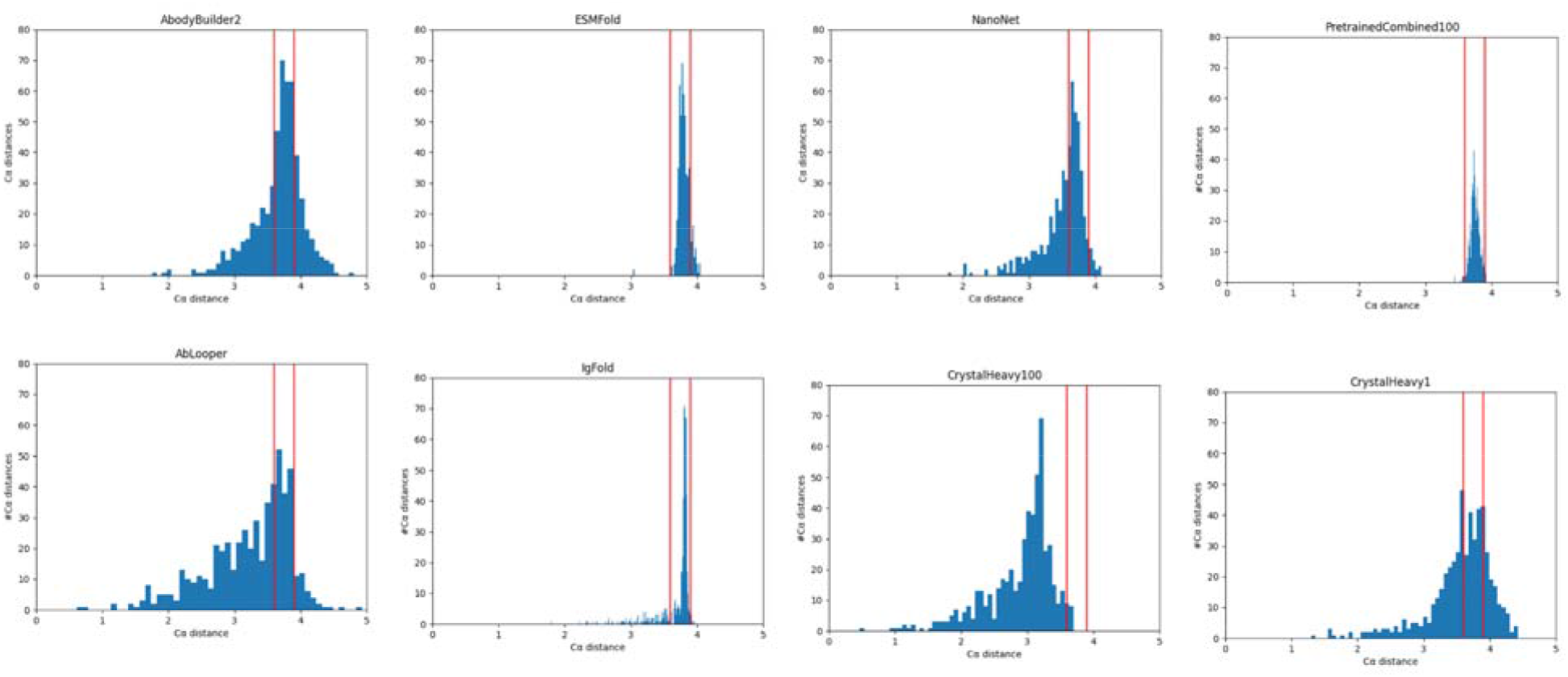
The distribution of Cα distances in the CDR-H3 predictions on the Rosetta test set. The red lines indicate the region of [3.6Å 3.9Å]. The distributions correspond to the predictions presented in Figure 3. Please note that the histograms have different bin sizes to fit the same axes in each image.

These results suggest that the pre-training strategy we introduced could have an effect on the number of correct peptide bonds, without notable detriment to CDR-H3 prediction accuracy with respect to other methods, despite simplistic model and training regimen.

### Effect of predicting out-of-distribution CDR-H3 structures

Antibody structural modeling tools need to be able to generalize to the vast number of conformations these molecules can adopt ^31^. Therefore, we checked to what extent different networks can produce physically plausible out-of-distribution sequences. For this purpose, we randomly generated ten CDR-H3s, between lengths [10-19] for each of the sequences in the Rosetta test set. This was supposed to be a the most extreme case of introducing the diversity, as opposed to following the natural substitution profiles ^32^ or sampling novel sequences from the OAS ^22,23^. It was checked that none of the generated CDR-H3s were in the crystal structure or the pre-training datasets. This resulted in 450 sequences.

We ran the predictions on our pre-trained model PretrainedCombined100, those without the benefit of pre-training, CrystalHeavy1 and CrystalHeavy100 as well as on IgFold, ESMFold, AbLooper and AbodyBuilder2. The AbodyBuilder2 predictions (final refined models) were used by AbLooper for remodeling the CDRs.

For each model, we calculated the number of bad Cα values (<3.6Å and >3.9Å). The number of CDR-H3 bad Cα distances for each model is given in Table 6. It is evident that in their unrefined form, IgFold, AbodyBuilder2 and AbLooper produce a large number of Cα-Cα distances falling outside of [3.6Å, 3.9Å] range. As on the proper Rosetta test-set, IgFold appears to produce a smaller amount of distances that would need to be refined as compared to AbLooper, AbodyBuilder2 or our not pre-trained models. From the previously developed methods, ESMFold does the best job of rapidly producing a low (509) number of incorrect bonds. Note that though the 509 values fall outside our range of [3.6Å, 3.9Å], the distribution is not too wide, not venturing below 2Å, which is not the case for other methods other than our pre-trained model (Figure 5). Note however that ESMFold achieves such performance without the benefit of costly refinement and produces full atom models, as opposed to simplistic Cα-only residual networks, trained here.

**Table 6.** Number of bad bonds in the artificial CDR-H3s. We generated 100 random CDR-H3s in the length range between 10-20 residues. Each of the structures was modeled by IgFold, AbodyBuilder2, ESMFold, NanoNet, our pre-trained model PretrainedCombyined100 and the not-pretrained varieties (CrystalHeavy100 and CrystalHeavy1). We report the number of Cα bonds that fall outside of the [3.6Å, 3.9Å] range for the CDR-H3.

| Model | #$C\alpha$-$C\alpha$ $< 3.6\text{\AA}$, $C\alpha$-$C\alpha$ $> 3.9\text{\AA}$ |
| --- | --- |
| PretrainedCombined100 | 517 |
| CrystalHeavy1 | 4,686 |
| CrystalHeavy100 | 5,837 |
| NanoNet | 4,005 |
| AbLooper | 5,035 |
| IgFold | 2,554 |
| AbodyBuilder2 | 4,097 |
| ESMFold | 509 |

**Figure 5.**
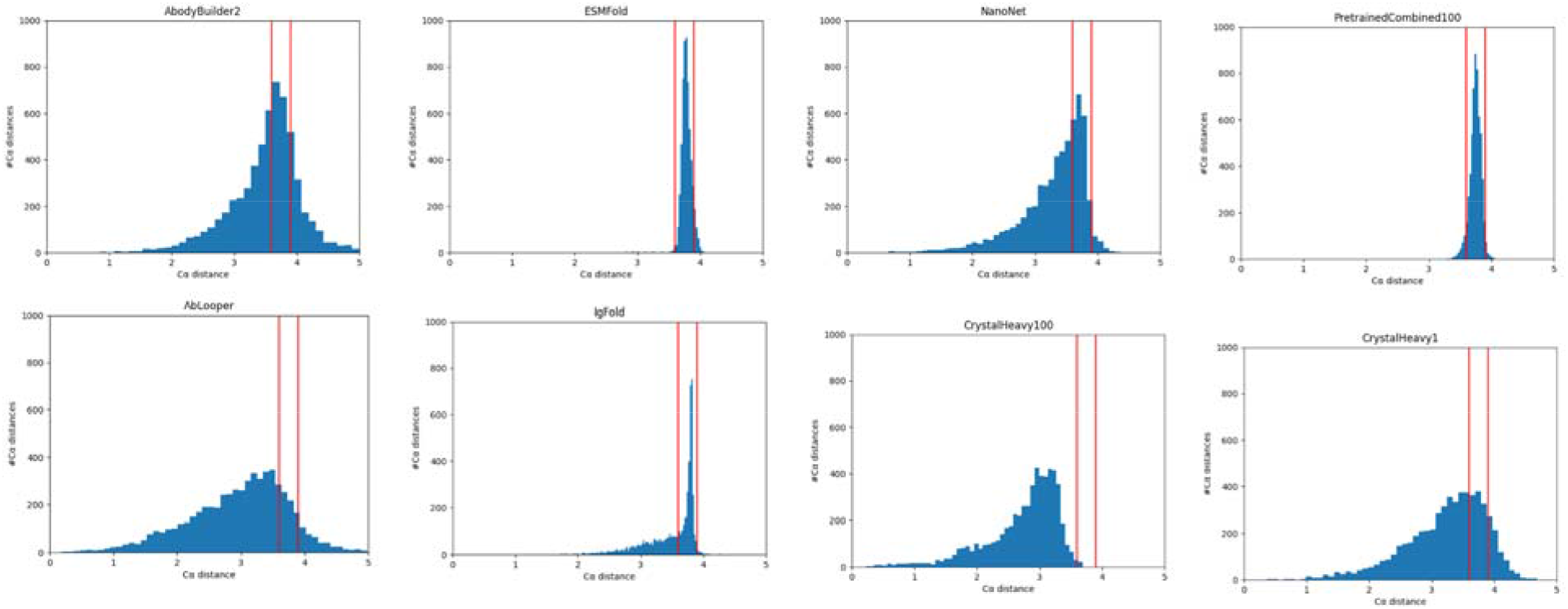
The distribution of Cα distances in the CDR-H3 predictions of the CDR-H3s randomized on the Rosetta test set. For each of the 45 structures in the Rosetta test set, we generated 10 random IMGT CDR-H3s in the length range [10 20] and we plot the histograms of the consecutive Cα distances in these. The red lines indicate the region of [3.6Å 3.9Å]. Please note that the histograms have different bins sizes to fit the same axes in each image.

Most importantly from the point of view of pre-training, CrystalHeavy100 model, which did not get the benefit of pre-training but a very large Cα parameter, produces the largest amount of bad distances, at 5,740. The model with a small Cα parameter but also without pre-training (CrystalHeavy1), also achieves a bad result in terms of number of peptide bonds (4,686). By contrast, the pre-trained model produces only 517 incorrect contacts. Since the only difference between the methods was the benefit of pretraining, we venture a statement that the network benefited from learning more correct physical bonds from a large pre-trained dataset that is able to generalize to physically plausible shapes of unseen (though perhaps unrealistic) CDR-H3s.

## Discussion

We addressed the issue of generating structural antibody models of better physical quality without detriment to prediction of the backbone Cα. We trained a very simplistic deep learning model with a rudimentary loss function to minimize the risk of better peptide bond distances being the result of ingenious model architectures ^16–18^. We demonstrated that even such simple models can learn better quality peptide bonds if given the benefit of pre-training on a large augmented set of refined antibody structures.

Producing good quality antibody models fast is crucial for practical applications of such methods in therapeutic development. During a phage-display or animal immunization campaigns, it is not uncommon to produce a number of sequences in the region of 10^5^-10^6^ or even 10^11 33^. Current methods such as IgFold, AbodyBuilder2, and ESMFold produce results of comparable quality in matter of seconds, which is an order of magnitude improvement versus the pioneer, AlphaFold2. However, even assuming an optimistic scenario that with refinement such methods would take only one second, 10^6^ sequences would still take approximately 11 days on a powerful GPU.

This could be a further hindrance when one wishes to study the entire space of antibodies, of which there is estimated to be 10^18^ unique molecules ^31^. A relatively tiny snapshot of this space in the form of Observed Antibody Space, currently holds 2,426,036,223 unique antibody sequences ^22,23^. Assuming an optimistic running time of 1s per model, covering all of these sequences on a powerful GPU would take approximately 673,898 hours or around 75 years. Producing physically accurate predictions without the need for energy refinement can reduce the running time, and thus make large-scale annotation of the antibody space within reach.

Nevertheless, the current efforts to model the available antibody space are not in vain as it was demonstrated that this could be done using even very rudimentary methods ^34^. We are employing such a natural dataset for pre-training obtained from public response structures. The authors of IgFold and AbodyBuilder2 also make available snapshots of more than 100,000 paired sequences available in OAS. Since snapshots provided by IgFold and AbodyBuilder2 use more advanced tools that we employed for pre-training here, we think that in the future they might provide an even better basis for structural pre-training. This raises the question of whether the structures in such pre-training datasets need to be only *‘*physically accurate*’* or be models that closely resemble the real structures that they model.

Furthermore, the proof of concept demonstrated here only took into account the peptide bond. It is not unreasonable to imagine that overall plausibility of the structure produced by the network might improve in other areas, such as side chains, the number of amino acids with D-stereochemistry or clashes ^24^. All such considerations hinge on presenting the network with a large, augmented dataset so it can develop a basis of inference of correct physical geometries that can be generalized to the vast space of antibody molecules.

As we demonstrated with the artificial CDR-H3 exercise, many networks show unstable behavior (non-physical peptide bond lengths) when predicting novel sequences, suggesting limited generalizability. Developing plausible novel antibodies is not only the domain of structure prediction but also generative modeling ^26,35^. While our approach of introducing augmented dataset for model pre-training improved model stability in CDR-H3 prediction, it might also be applicable to producing novel physically plausible structures using generative modeling regimes.

Altogether, employing a large, augmented dataset for structural pre-training appears to produce physically plausible predictions for known structures as well as randomly generated ones. Therefore we hope that our presented strategy for structural pre-training would benefit the many antibody structural modeling tools available currently.

## Methods

### Datasets

#### Rosetta test set

We employed the *‘*Rosetta*’* test set introduced previously as multiple methods employed it subsequently for benchmarking. The original Rosetta test set consisted of 48 Fvs. We removed three of them because of low quality, for instance incomplete chains. The final dataset consisted of the following 45 PDBs : 3gnm, 2xwt, 2vxv, 3i9g, 2d7t, 2ypv, 1mlb, 2w60, 1gig, 3g5y, 2adf, 3liz, 3p0y, 3hnt, 3e8u, 3mxw, 2fbj, 1nlb, 3go1, 2fb4, 3mlr, 1jfq, 3nps, 2e27, 1fns, 2v17, 1seq, 4nzu, 1mqk, 4f57, 4hpy, 2r8s, 1mfa, 3oz9, 3hc4, 1jpt, 3t65, 1oaq, 3v0w, 3lmj, 3m8o, 4h20, 1dlf, 3eo9, 3giz.

#### Crystal datasets

We downloaded the antibodies from the Protein Data Bank on March 13^th^ 2022. We removed the structures with identical CDR-H3 or CDR-L3 loops. Only X-ray structures with resolution better than 3Å were left in. We removed all the structures that had the same CDR-H3 or CDR-L3 as in the Rosetta test set. The final dataset consisted of 1,628 training and 185 validation heavy chain structures that we term as CrystalHeavy. The CrystalLight dataset consisted of 1,302 training and 149 validation light chain structures. The CrystalCombined dataset was created by taking the intersection of the two datasets resulting in 2966 training and 298 validation structures.

#### Pre-training datasets

We employed two pre-training datasets *‘*Natural*’* and *‘*Homology*’*. The Natural dataset was created externally as the sample of the public response structures ^30^. We created the Homology dataset by reproducing the homology modeling protocol introduced by AbodyBuilder ^20^ via: 1) selecting the closest template framework from the PDB ^34^, 2) orientating the chains by closest matching pair 3) modeling the loops by FREAD ^36^ and refining the predictions by PEARS ^37^. In both the Natural and Homology datasets, we removed all the structures where CDR-H3 or CDR-L3 matched this in the CrystalHeavy or CrystalLight datasets. We further removed structures that had more than 1Å RMSD to CDR-H3 or CDR-L3 in the Rosetta test set. This resulted in 18,959 training, 2369 validation and 2369 test structures in the natural dataset. Likewise, the Homology dataset consisted of 37,672 training, 4709 validation and 4709 test structures. Since we demonstrated that combining heavy and light chains for prediction appears to improve the peptide bond length without detriment to shape prediction quality we created a PretrainCombined dataset by merging the heavy and light chains in the Homology and Natural datasets. This resulted in 113,262 structures for training 11,326 for validation and 11,326 for test.

### Models

#### Model for pre-training

We employed the simplest deep learning architecture we could identify that reported heavy chain modeling of antibodies, which was NanoNet ^14^. In brief it consists of seven convolutional residual blocks. It has 1.9m parameters which puts it on the lower spectrum with respect to models trained on the basis of 15B parameters language models (ESMFold) or on the basis of AlPhaFold (AbodyBuilder2). The lightweight architecture ensured that we could do multiple iterations and variations of the models to test the effects of the structural pre-training strategy (e.g. AbodyBuilder achieves very good results but it takes several weeks to train on a powerful GPU). Furthermore, the lightweight architecture reduced the risk that any effects we might observe of better peptide bond distances, would be accounted for by ingenious architectures or training regimes. The method does not come with a training loop so the architecture and training functions had to be reimplemented from scratch. For the training loss we employ the Mean Squared Error together with a loss on the consecutive distances of Cα. The Cα loss was parameterized with weight, biasing the predictions towards putting more attention to Cα distances.

#### Models for benchmarking

We employed five models for benchmarking, AbodyBuilder2, AbLooper, IgFold, ESMFold and NanoNet. The choice was motivated by the availability of the methods and the fact that they are state-of-the art. The models coming from AbodyBuilder2, AbLooper and IgFold did not undergo refinement, which is the full protocol in each case. This was to compare the results of predictions coming straight from the network. All the methods are freely available and they were obtained from the repositories advertised in the corresponding publications.

### Quality Metrics

#### Cα-Cα peptide bond distances

We calculated the Cα-Cα distances between the consecutive atoms as the simple euclidean distance of the coordinates. Our datasets were not allowed to have structures with missing amino acids, resulting in incorrectly large Cα-Cα distances as a result of a missing residue or a stretch of these in between. All the Cα-Cα distances that were falling outside of the [3.6 3.9] range we considered as incorrect, which was an arbitrary choice based on observation of the distance in the crystal structures.

#### RMSD

Whenever we report the Root Mean Square Deviation between prediction and native structures we only take Cα into account. The RMSDs for the H3s were calculated in two modes - by Fv alignment (Employed in Table 3) and by CDR-H3 alignment (employed in Figure 3). In the Fv alignment, the Cα coordinates of the entire variable region are aligned using the KABSCH algorithm in the python *rmsd* module and the RMSD is calculated on the IMGT-defined CDR-H3 only. In the CDR-H3 alignment, the CDR-H3 Cα only are aligned and the RMSD calculated. Clearly, the CDR-H3 alignment should typically produce lower numbers.

